# Galvanotactic directionality of cell groups depends on group size

**DOI:** 10.1101/2024.08.13.607794

**Authors:** Calina Copos, Yao-Hui Sun, Kan Zhu, Yan Zhang, Brian Reid, Bruce Draper, Francis Lin, Haicen Yue, Yelena Bernadskaya, Min Zhao, Alex Mogilner

**Affiliations:** Department of Biology and Department of Mathematics, Northeastern University, Boston, MA 02115; Department of Ophthalmology and Vision Science and Department of Dermatology, School of Medicine, University of California, Davis, Sacramento, CA 95817; Department of Occupational and Environmental Health, Hangzhou Normal University School of Public Health, Hangzhou 311121, China; Department of Molecular and Cellular Biology, University of California, Davis, Davis, CA 95616; Department of Physics and Astronomy, University of Manitoba, Winnipeg, MB R3T 2N2, Canada; Department of Physics, University of Vermont, Burlington, VT 05405; Courant Institute and Department of Biology, New York University, New York, NY 10012

## Abstract

Motile cells migrate directionally in the electric field in a process known as galvanotaxis, important and under-investigated phenomenon in wound healing and development. We previously reported that individual fish keratocyte cells migrate to the cathode in electric fields, that inhibition of PI3 kinase reverses single cells to the anode, and that large cohesive groups of either unperturbed or PI3K-inhibited cells migrate to the cathode. Here we find that small uninhibited cell groups move to the cathode, while small groups of PI3K-inhibited cells move to the anode. Small groups move faster than large groups, and groups of unperturbed cells move faster than PI3K-inhibited cell groups of comparable sizes. Shapes and sizes of large groups change little when they start migrating, while size and shapes of small groups change significantly, lamellipodia disappear from the rear edges of these groups, and their shapes start to resemble giant single cells. Our results are consistent with the computational model, according to which cells inside and at the edge of the groups pool their propulsive forces to move but interpret directional signals differently. Namely, cells in the group interior are directed to the cathode independently of their chemical state. Meanwhile, the edge cells behave like individual cells: they are directed to the cathode/anode in uninhibited/PI3K-inhibited groups, respectively. As a result, all cells drive uninhibited groups to the cathode, while larger PI3K-inhibited groups are directed by cell majority in the group interior to the cathode, while majority of the edge cells in small groups win the tug-of-war driving these groups to the anode.

**Significance statement:** Motile cells migrate directionally in electric fields. This behavior – galvanotaxis – is important in many physiological phenomena. Individual fish keratocytes migrate to the cathode, while inhibition of PI3K reverses single cells to the anode. Uninhibited cell groups move to the cathode. Surprisingly, groups of PI3K-inhibited cells exhibit bidirectional behavior: larger/smaller groups move to the cathode/anode, respectively. A mechanical model suggests that inner and outer cells interpret directional signals differently, and that a tug-of-war between the outer and inner cells directs the cell groups. These results shed light on general principles of collective cell migration.

## INTRODUCTION

Cells migrate collectively, as cohesive groups, in development, wound healing, and tumor invasion [1-3]. Understanding the coordinated movement and directionality of these groups is an important open problem. Experimental research [1, 2, 4] on and modeling [5-8] of cell groups migrating spontaneously or in chemical gradients brought much insight into mechanics and directional cell behavior. A few conceptual models emerged from this research: A) Leader cells [9, 10] are polarized and actively migrating in the direction of an external cue. The remaining cells follow the leader cells passively [7]. B) Inner cells in the group polarize in the direction of the external cue and migrate actively, while cells at the group edge do not respond to the directional signal and are dragged and pushed along by the inner cells [8, 11]. C) The group is integrated mechanically [12, 13], so that all cells are tightly interlinked into a supracellular tissue by cytoskeletal structures spanning the whole group [14], and the groups migrates like one giant cell. More complex and nuanced models have also been considered [15, 16].

Much research on collective cell migration was done on groups moving in chemical gradients, yet there are other directional cues that cells encounter. Cells are electrical units: they transport ions across membranes, and as a result are surrounded and regulated by electrical fields and currents [3, 12]. Ability of cells to sense the direction of an electric field (EF) – galvanotaxis – plays important roles in development [17], wound healing [18], and regeneration [19]. Some types of cells (i.e., keratinocytes) individually migrate to the cathode in EF, others (i.e., fibroblasts) – to the anode [20]. Respective galvanotactic signals may be as potent as, or even more important than, chemotactic signals [21]. Electrically sensitive cells is the rule, not the exception [21]. Most of research on galvanotaxis was done on single cells, but recent studies also investigate collective migration in EF of epithelial cell sheets [22, 23], MDCK cell groups and keratinocytes [11, 24], corneal epithelial cells [25], and HaCaT cellular monolayers [26].

One cell type, fish epidermal keratocyte, has been instrumental in studying mechanisms of galvanotaxis due to its fast and steady locomotion, simple shape, and well-understood motile mechanics [27]. Keratocytes sense EF and move to the cathode [28]. Physically, keratocytes likely sense EF by harnessing electrophoresis of charged mobile transmembrane proteins, which aggregate to one of the cell’s sides in EF [29] and serve either as receptors activating intracellular signaling relays, or as scaffolds for such receptors. Inhibition of PI3-Kinase (PI3K, for brevity) redirects these cells to the anode [29, 30]. Both single keratocyte migration, and spontaneous migration of groups of zebrafish [31] and gold fish [5] keratocytes, as well as EF-induced collective movements of zebrafish [32, 33] and gold fish [28] keratocytes have been characterized.

PI3K is one of the key signaling molecules in pathways transducing the electrical signal to the cell motility apparatus [34], and individual PI3K-inhibited cells migrate to the anode (positive electric terminal), with large lamellipodia leading the way [30], oppositely to unperturbed individual cell migrating to the cathode (negative terminal). In [33], we reported that very large zebrafish keratocyte groups of thousands of uninhibited or PI3K-inhibited cells migrate to the cathode. To understand why individual PI3K-inhibited cells migrate to the anode but large groups of such cells – to the cathode, in this study, we use groups of several to tens to hundreds of cells to investigate their polarization, migration and directional sensing in EF. We find that while larger (hundreds of PI3K-inhibited cells) groups move to the cathode, small (tens of PI3K-inhibited cells) groups move to the anode. These behaviors are consistent with a model according to which cells at the group edge sense EF in a similar way to the individual cells and tend to go to the cathode or anode depending on their chemical state (unperturbed or PI3K-inhibited, respectively). Meanwhile, cells in the group’s interior are not passive followers, but rather tend to polarize and move to the cathode, independently of their chemical state. In unperturbed groups, both inner and outer cells are driven to the cathode, and the whole group moves to the cathode. A tug-of-war between the inner and outer cells in PI3K-inhibited groups determines the group’s directionality: in larger groups, the inner cell majority drives the group to the cathode; in small groups, the outer cell majority drives the group to the anode.

## RESULTS

### Collective behavior of two cohesive cells suggests directional / mechanical rules for outer cells

To understand directionality of a cohesive pair of cells in EF, we observed such PI3K-inhibited pair (Fig. 1A; we could not find a two-unperturbed-cell group) and saw that before EF was on, each of the two cells resembled a half-disc, with shared segment of a boundary and semi-circular protruding lamellipodium. Straightforward interpretation of this morphology is that the common cohesive rear of this pair does not protrude due to the contact inhibition of locomotion [35], and that as far as this common rear position is determined by the cohesive intercellular contact, both cells become strongly polarized outward due to the ubiquitous tendency of motile cells to generate oppositely oriented contractile rear and protrusive front [36, 37]. The clear indication of the propulsive force trying to move each cell away from the shared rear is that sometime the cohesion in the central parts of the rear ruptures, and the cells’ rears, held together just by two contacts at the sides, curve outward pulled by respective protrusions (frame 2 of Fig. 1A). When the EF is on (blue-bracketed frames of Fig. 1A), initially the cells stay cohesive, but, notably, start shifting together slowly in the anode direction (Movie 1 and compare two first blue-bracketed frames of Fig. 1A). This can be interpreted as the anode-facing lamellipodium of one of the cells being propelled by a stronger force than the oppositely cathode-facing lamellipodium of another cell, suggesting that there is a superposition of two types of forces. One is a constant-amplitude outward force trying to propel the cell in the direction normal to its rear, orientation of which is geometrically imposed by the contact with inner cells in the group, and another, EF-induced, directed along the anode-cathode axis, with amplitude graded by the angle between the anode (or cathode) and angle between this axis and the geometrically constrained rear-front axis of the lamellipodium. We use this concept to build the model of cell group collective directionality below.

**Figure 1.**
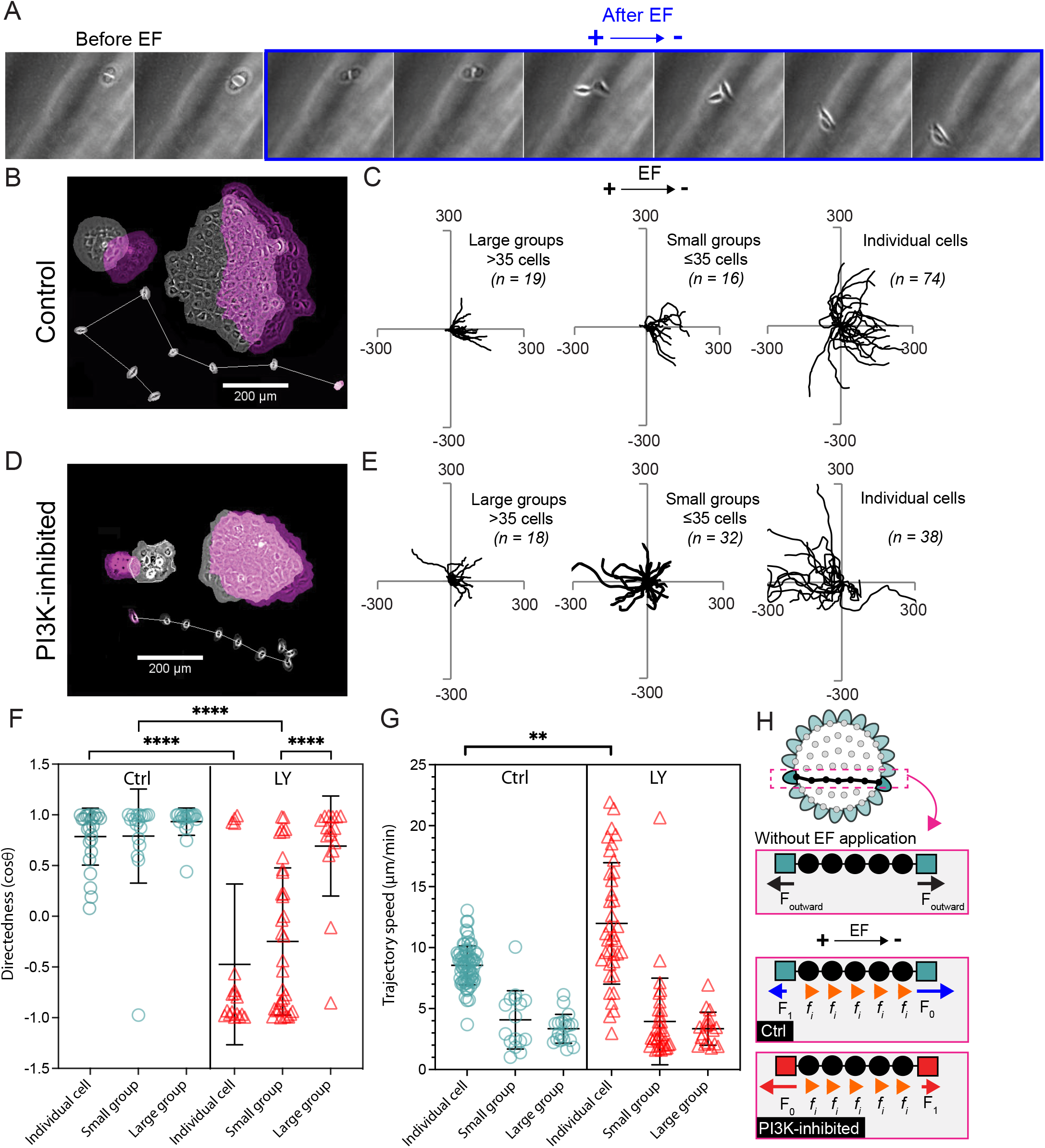
EF guides individual keratocytes and groups to the cathode but directs individual cells and small groups to the anode in the presence of the PI3K inhibitor. (A) Snapshots of a pair of PI3K-inhibited cells imaged before and during 30 minutes after 1 V/cm EF application (see also Movie 1; cathode is at the right). (B and D) Overlays of representative large and small cell groups and of one individual cell (with its migratory trajectory) before (gray) and 2 hours after (magenta) 1 V/cm EF application in (B) control and (D) PI3K inhibition. (C and E) Migratory trajectories of centroids of large (>35 and <400 cells), small (≤35 cells) cell groups and individual keratocytes after applying 1 V/cm EF for 30 minutes in (C) control and (E) PI3K inhibition. The distances along x and y axes are in µm. The tracks are re-centered to the same starting point at the moment of EF application; trajectories are shown over 30 minutes after EF application. (F and G) Comparison of the directedness (cosine of angle between the anode-cathode vector and the vector connecting the beginning (at the moment of EF application) and end (30 minutes after EF application) positions of the cell or group centroid; see Methods) and of the trajectory speed (speed of the centroid averaged over 30 min in EF, in µm /min) of the of individual cells, small and large groups, trajectories of which are shown in C and E. The quantification is done over a 30-minute period post-EF application. Horizontal lines indicate distribution mean with standard deviation error bars. ****p<2.7e-6; **p<2e-3; ns, not significant (p>0.05), one-way analysis of variance (two-tailed t-test). (H) One-dimensional theoretical model of propulsive forces in collective galvanotaxis.

After a few minutes of the tug-of-war between the two cells, they break contact, the initially cathode-facing cell makes a pivot to the anode and becomes a follower cell moving behind, and in touch with, the leader (last four frames of Fig. 1A). This hints at the phenomenon described below: when the group starts following the ‘correct’, EF-informed, direction, the lamellipodia at the group’s rear can destabilize, and the group’s rear is made of rears of outer cells on that side.

### Directionality of EF-guided cells and cell groups depends on the cell chemical state and group size

To confirm that the two-cell example is not an exception and that small groups of PI3K-inhibited cells move to the anode, we investigated keratocyte groups moving in physiological-range EF of magnitude ∼1 V/cm [38], and recorded 30-min long trajectories of single cells and groups of various sizes (Fig. 1B-E). In the case of unperturbed cells, both single cells and groups of all sizes migrated to the cathode (Fig. 1B,C). In the case of PI3K-inhibited cells, while large (more than 35 cells) groups also migrated to the cathode, single cells and small (less than 35 cells) groups migrated oppositely, to the anode (Fig. 1D,E). Statistics of the directedness – cosine of the angle between the vector connecting the initial and final positions of the groups’ centroids and the cathode direction – confirm this result quantitatively (Fig. 1F). We also measured trajectory speed of the cells and groups (Fig. 1G) and found that individual cells, both unperturbed and PI3K-inhibited, moved faster than respective groups, and that the small groups, despite large variations of the speed between the groups, moved faster than the large groups (Fig. 1G; small and large unperturbed groups moved with speeds ≈ 4.07 μm/min and ≈ 3.34 μm/min, respectively, while small and large PI3K-inhibited groups moved with speeds ≈ 3.94 μm/min and ≈ 3.35 μm/min, respectively).

### 1D mechanical model of the tug-of-war between the outer and inner cells explains the size-dependent group directionality

Quantitative modeling has been instrumental in understanding collective cell directionality in chemical gradients [7, 39, 40], and individual cell directionality in EF [41, 42]. Individual [43-46] and collective [13, 43, 47] cell mechanics have also been modeled, and altogether these theories suggest the following simple conceptual model that helps to begin understanding the bidirectional group motility in EF (Fig. 1H). Let us consider a one dimensional (1D) cohesive chain of cells stretching from the anodal to cathodal edge of the group (Fig. 1H), and for now, consider this cell chain in isolation, without contacts with other cells on the sides. Based on measurements of responses of the cells to EF within very large groups [33], we will assume that all cells in the chain, including the inner and outer (edge) cells, sense EF and respond to it individually and independently. Following previous modeling work, we will also assume that each cell in the chain responds to EF by generating a propulsion force, direction and magnitude of which depends on the cell position within the group.

As suggested by the 2-cell behavior described above, without EF, the outer cells generate outward propulsive forces of equal magnitude, *F*_*outward*_ (Fig. 1H). It would be natural to assume that the inner cells are not directional without EF, and then two outer cells in the 1D chain balance each other and the cohesive group is stationary. Based on the 2-PI3K-inhibited-cell behavior such, the magnitude of the anode-directed force of the anode-facing leader (left outer) cell, −*F*_0_, is greater than the magnitude of the cathode-directed force of the trailer (right outer) cell, *F*_1_ (Fig. 1H). (Positive/negative force is directed to the cathode on the right / anode on the left, respectively.) If in the presence of EF the inner cells are not directional, then the cell chain of any size would then experience the net anode-directed propulsive force (−*F*_0_ + *F*_1_) and move to anode, contradicting the data. Thus, we posit that each inner cell generates a cathode-directed propulsion force + *f*_*i*_ in any chemical state. Similarly, it is natural to assume that in the chain of unperturbed cells, while each inner cell generates propulsion force + *f*_*i*_, the leader cell is on the cathodal side and exerts the highest force, *F*_0_, to the right, and the trailer cell is on the anodal side and exerts the lower force, −*F*_1_, to the left. In principle, the forces’ magnitudes could be different in the unperturbed and PI3K-inhibited cases, but for simplicity we will keep them the same. Similarly, the inner cell propulsive forces can depend on the position inside the group [13], but for simplicity we will keep them position independent. Several experimental studies indicated that leader (outer) cells exert higher forces than follower (inner) cells [4, 13, 48, 49], so we assume that *F*_0_ > *F*_1_ > *f*_*i*_. If there are *N* inner cells in the chain, then the total force propelling the group is equal to (*F*_0_ − *F*_1_ + *Nf*_*i*_) in the unperturbed case, and to (*F*_1_ − *F*_0_ + *Nf*_*i*_) in the PI3K-inhibited case.

This simple 1D model makes several predictions. First, both unperturbed and PI3K-inhibited *large* groups are propelled with force approximately equal to *Nf*_*i*_ (if *N* >> 1, then (*F*_0_ − *F*_1_ + *Nf*_*i*_) ≈ (*F*_1_ − *F*_0_ + *Nf*_*i*_) ≈ *Nf*_*i*_). It is then logical to conclude that unperturbed and PI3K-inhibited *large* groups move with the same speed. Second, we know that large groups in both chemical states move to the cathode, indicating that *f*_*i*_ > 0 in both cases. Thus, inner cells respond to EF by polarizing to the cathode in both chemical states. Third, small unperturbed groups should move to the cathode, because if *F*_0_ > *F*_1_and *f*_*i*_ > 0,then (*F*_0_ − *F*_1_ + *Nf*_*i*_) > 0 for any *N*. However, small PI3K-inhibited groups should move to the anode, because if *F*_0_ > *F*_1_, then (*F*_1_ − *F*_0_ + *Nf*_*i*_) < 0 for *N* = 0 and for small *N* < (*F*_0_ − *F*_1_) / *f*_*i*_. Fourth, small unperturbed groups should move faster than small PI3K-inhibited groups of the same size because |*F*_0_ − *F*_1_ + *Nf*_*i*_ |> | *F*_1_ − *F*_0_ + *Nf*_*i*_ |if *F*_0_ > *F*_1_(simply speaking, because in unperturbed groups, inner and outer cells pull the group in the same, cathodal, direction, but in PI3K-inhibited groups, inner cells pull the group in the cathodal direction, while outer cells altogether push the group in the opposite direction). The predictions about equal speeds of large unperturbed and PI3K-inhibited groups and of speed of small unperturbed group being greater than speed of small PI3K-inhibited group agree with the measurements. Full mathematical analysis of the 1D model is presented in the SI and in Fig. S2.

These predictions should be qualitatively valid in the realistic 2D case, too. For roundish groups of *N* cells, the number of outer cells scales as 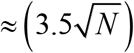, while the number of inner cells scales as 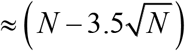.Large groups, both unperturbed and PI3K-inhibited, move to the cathode, because in such groups inner cells are in majority 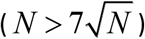 and win in the directional tug-of-war, while small groups 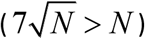 are driven by the outer cell majority, which is either cathodal or anodal directed depending on whether the group is unperturbed or PI3K-inhibited, respectively. It is interesting to note that the crossover from the inner to outer cell majority corresponds to the group size *N* ≈ 50 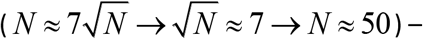 the same order of magnitude as the threshold size at which we observe the directionality switch (between 20 and 60-cell groups).

### Large groups do not change size and shape upon motility initiation, while smaller groups polarize and lose lamellipodia at the rear boundaries

To investigate the shapes and sizes of the galvanotactic cell groups, we excluded the groups which appeared polarized and moving directionally before EF was on and measured displacements, areas and aspect ratios, from 30 minutes before EF was on to 30 minutes after EF was on, of such small (6-35 cells) and large (37-200 cells) groups. Respective representative examples are shown in Fig. 2A-D and Movies 2-5, and statistics quantifying the EF-induced movements and shape changes can be gleaned from Fig. 2I.

**Figure 2.**
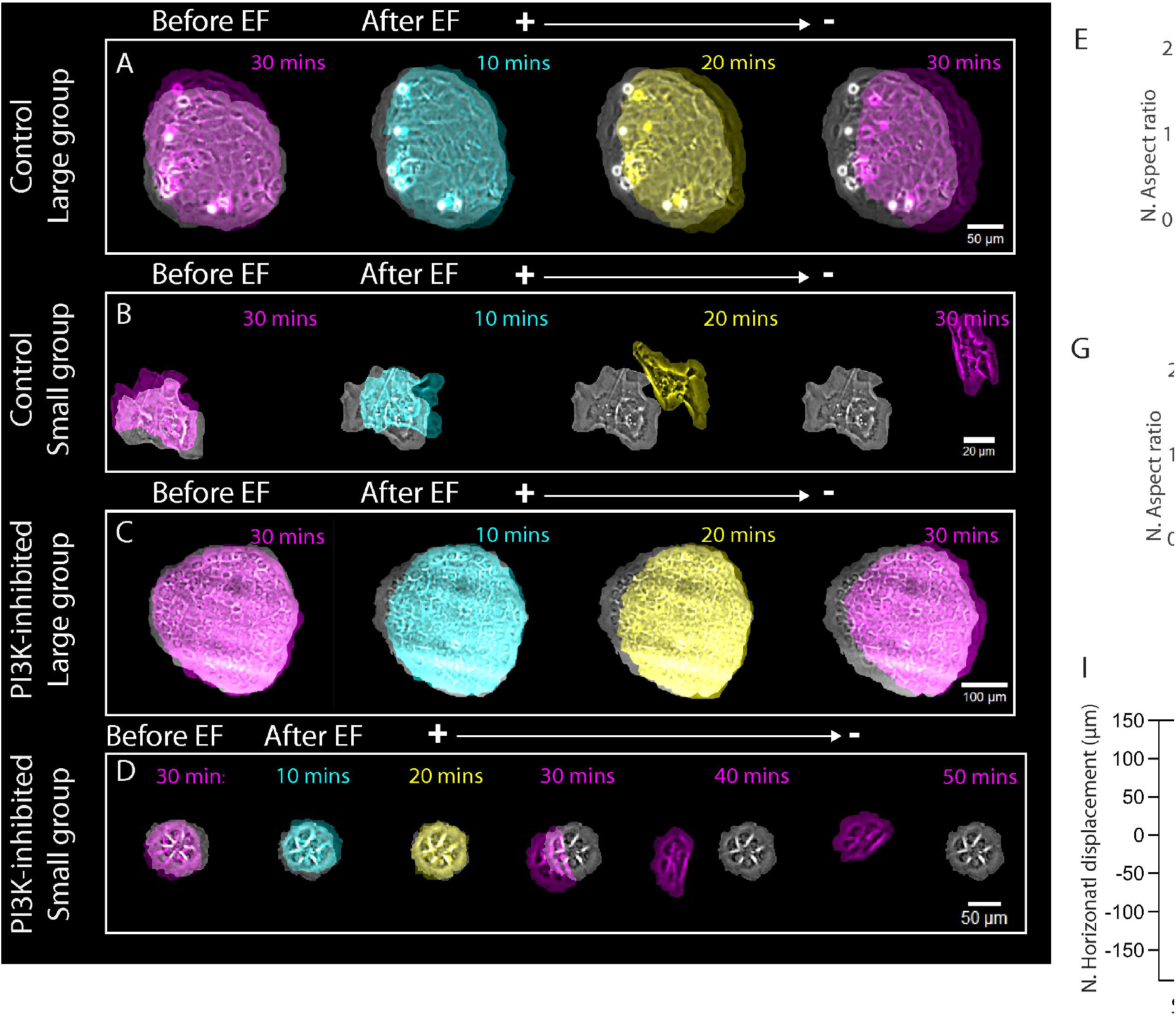
Significant morphological changes observed in small groups but not in larger groups. (A, B, C, and D) Overlays of representative large and small cell groups earlier (gray) and later (color) in control and PI3 inhibited cases. A 1 V/cm EF is applied with the cathode to the right. See also Movies 2-5. (E, F, G, and H) Normalized (by its value before EF application) aspect ratio and displacement along the EF axis over time in both large (E,G) and small (F,H) groups in (E, F) control and (G, H) PI3K-inhibited cases. 1 V/cm EF is applied at the 30-minute mark. (I) Comparisons of displacement, area, and aspect ratio changes induced by electric field in the absence or presence of the PI3K inhibitor (LY). Displacement along the EF axis is reported 30 minutes after 1 V/cm EF application in large and small cells groups in the absence (Ctrl) and presence (LY) of the PI3K inhibitor. Absolute values of the normalized area and aspect ratio changes are reported from right before to 30 minutes after 1 V/cm EF application. **p<3.6e-5; ns, not significant (p>0.05), one-way analysis of variance (two-tailed t-test). *p<0.04; ns, not significant (p>0.05), one-way analysis of variance (two-tailed t-test). Small control groups: n = 6, large control groups: n =7, small LY groups: n = 12, and large LY groups n = 5. Horizontal lines inside the boxes indicate distribution median, while the plus symbols show the mean. Tops and bottoms of each box indicate 75% (q_3_) and 25% (q_1_) percentiles, respectively. The whiskers extend to the highest and lowest values.

Large cell groups of both unperturbed and PI3K-inhibited cells moved to cathode (Fig. 2A,C,E,G), almost without changing shape (Fig. 2E,G,I). Areas of these groups also changed relatively little (Fig. 2I). Note that the displacements of both large unperturbed and PI3K-inhibited groups were very similar to each other, as the simple 1D model predicted. Notably, most outer cells had lamellipodia protruding outward all around the group boundary, including the groups’ rear (Fig. 2A,C, Movies 4,5 and Fig. S3).

Absolute values of the displacements of small unperturbed cell groups were greater than those of the small PI3K-inhibited groups, just as the simple 1D model predicted (Fig. 2I). Most notably, the aspect ratio of the small groups of both kinds changed significantly in EF – the groups became less roundish, shorter from rear to front and longer side-to-side (Fig. 2B,D,F,H and Fig. 2I). The groups’ shape (Fig. 2B,D), in fact, started to resemble the typical single motile cell shape. The reason for that was that most of the outer cells’ lamellipodia at the rears of the groups were lost (Fig. 2B,D, Movies 2,3), allowing the groups to pull its rear forward and inward, while the outer cells at the front and sides of the groups kept the lamellipodia stretching the group from side to side. The areas of the small groups decreased upon motility initiation going down more significantly than that of the large groups (Fig. 2I).

### 2D mechanical model with velocity-dependent slippery lamellipodial clutch explains the data

To see if the mechanical tug-of-war model could account for the data quantitatively and explain not only displacements but also shape changes of the motile groups, we turned to the 2D computational model (Fig. 3A). In the experiments, we observed that at the time scale of tens of minutes, the order of cells within the groups remained fixed – individual cells did not exchange neighbors, thus, the cells in the groups were cohesive with their neighbors, and so we modeled the mechanical links between neighboring cells as elastic. Each cell was assumed to resist its propulsion force by generating viscous-like resistance [13] characterized by the effective drag coefficient, same for all cells. Without EF, inner cells in the model did not generate any propulsive force, while the outer cells generated equal-magnitude protrusive forces directed outward perpendicularly to the local group edge (Fig. 3A) (as in [7, 39]). With EF, each inner cell in the model generated a constant cathode-oriented propulsion force independent of the chemical state, while the outer cells generated the geometry- and chemical state-dependent propulsion forces, as follows. Outer cells, lamellipodia of which faced the cathode, generated a cathode-oriented force *F*_0_ (unperturbed) or *F*_1_ (unperturbed) (Fig. 4A). Outer cells, lamellipodia of which faced the anode, generated an anode-oriented force *F*_1_ (unperturbed) or *F*_0_ (unperturbed). Lastly, we introduced the ‘slippery clutch’ mechanism, which is similar to models in [43, 46] and which is suggested by our observations: when the group moves directionally and the lamellipodia of the outer cells are dragged by the group against the direction, in which these lamellipodia try to protrude, these lamellipodia get destabilized, if the speed of the movement in the ‘wrong’ direction exceeds the threshold.

**Figure 3.**
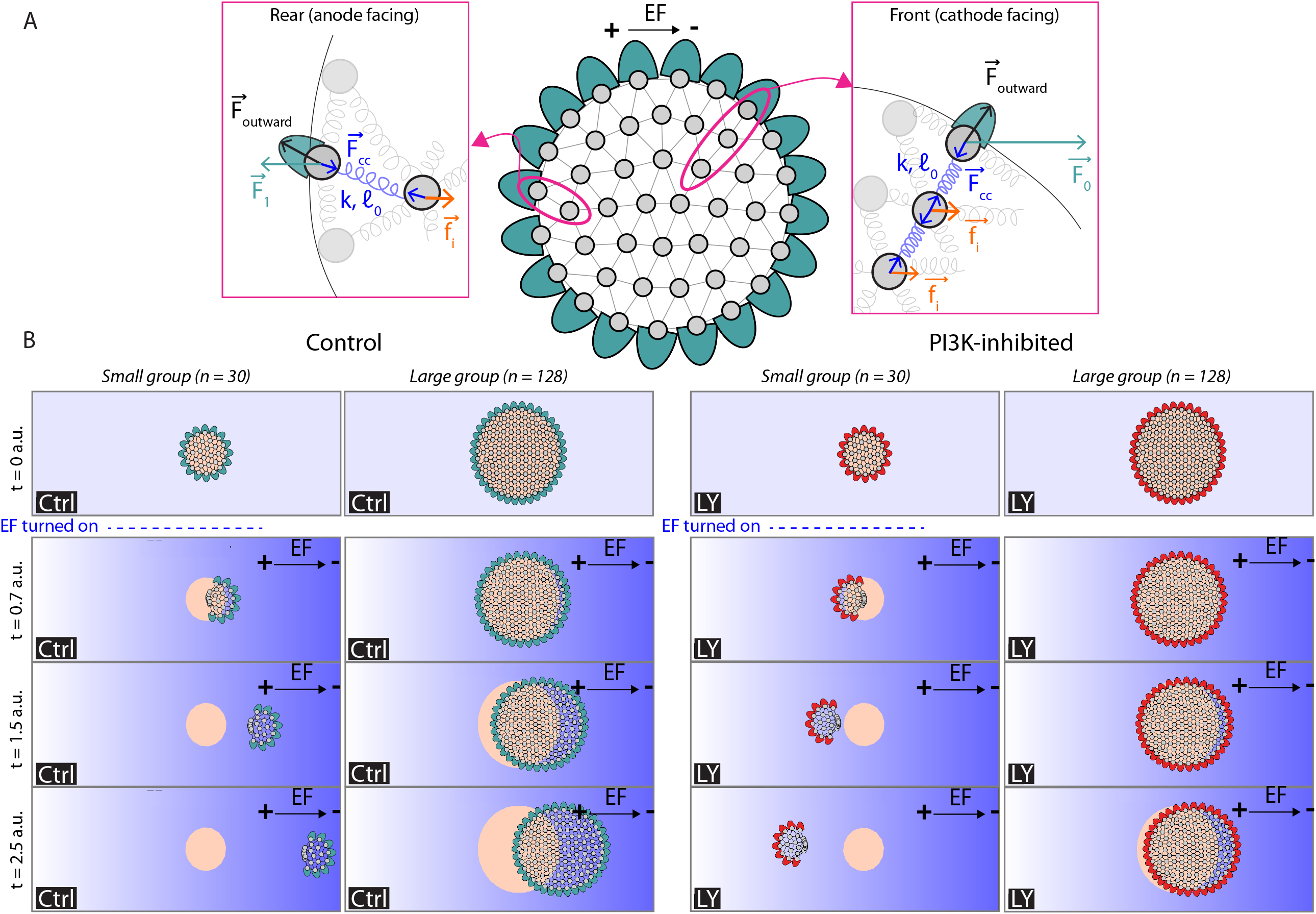
2D model with propulsion forces that depend on the cell position within the group and on the chemical state. (A) Diagram illustrating all components in the model, including cell-cell forces, EF-induced propulsion forces, and outward protrusive forces for outer cells in the outward normal direction. The cell-cell interaction is modeled with an elastic spring with nonzero resting length and stiffness. For unperturbed (Ctrl) groups, the outer cells (cyan) sense EF leading them to the cathode, and the propulsion forces at the front of the group are larger than those at the back (blue arrows). The inner cells generate propulsion forces pointing to the cathode (orange arrows). For the PI3K-inhibited group, the propulsion forces of the outer cells pointing to the anode are larger than those at the front (red arrows), while the active forces of inner cells are the same as in the unperturbed group (orange cells). (B) Simulation snapshots of the model at four distinct timepoints: initially (t = 0 a.u.) and after application of the EF at t = 0.5 a.u. (t = 0.7, 1.5, and 2.5 a.u) (see also movie 6). Cathode is always to the right. The initial configuration is shown in an orange overlay, while the later configuration is shown with individual cells (gray) and lamellipodia of engaged outer cells (cyan in control groups and red in PI3K-inhibited groups).

**Figure 4.**
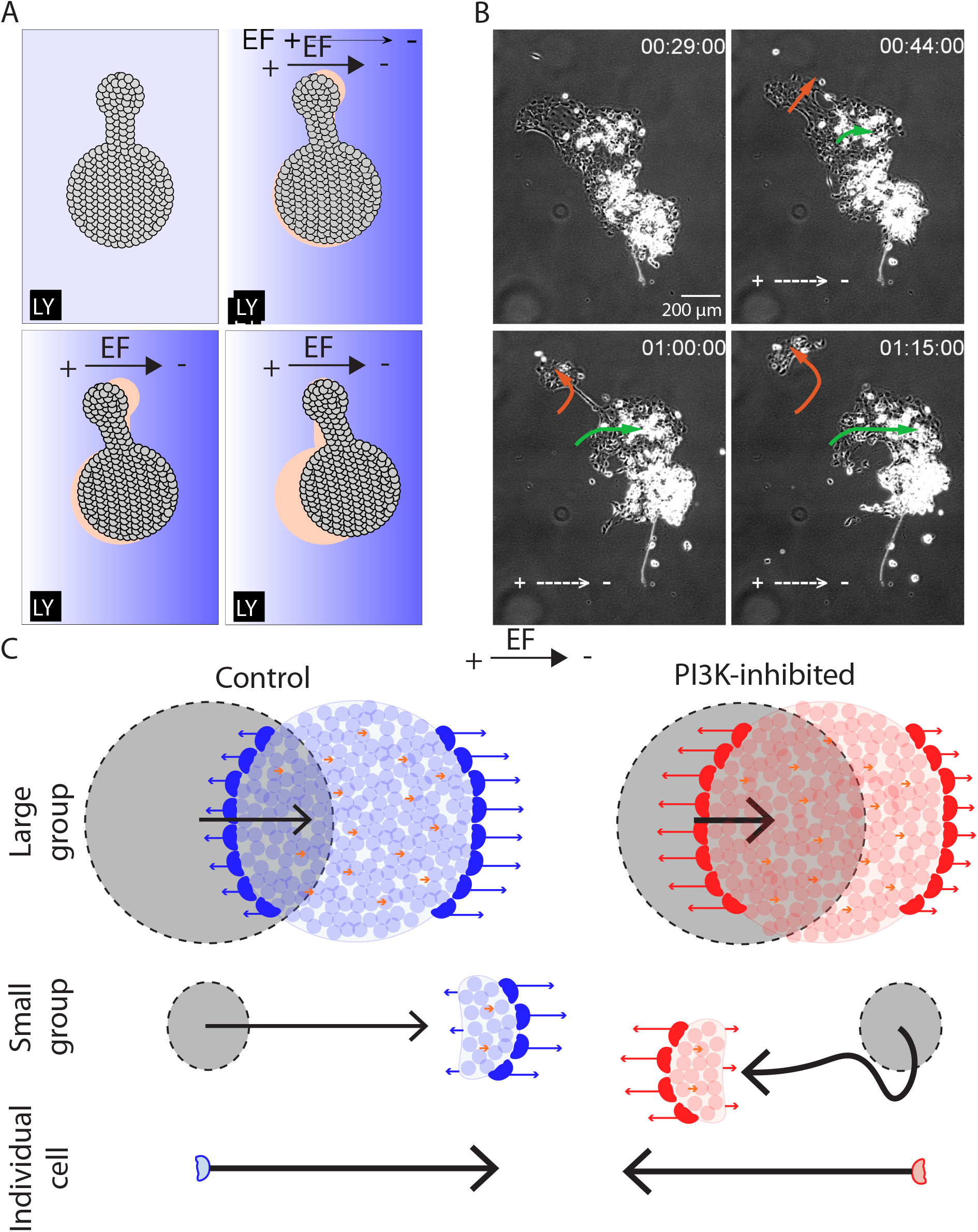
In the presence of 100 µM of the PI3K inhibitor, small group of cells splits off from a larger cell cluster. (A) Simulation snapshots of a dumbbell-shaped cluster of cells with n = 270. The orange background overlay shows the initial configuration. In the simulation, the cell-cell links, modeled as elastic bonds, cannot rupture (see also Movie 7). (B) Snapshots of a large group of cells with a 1 V/cm EF in the presence of 100 µM of the PI3K inhibitor compound (see also Movie 8). Arrows indicate the displacement of the original group (green) and the small group (orange) that eventually splits off. The EF is applied at the 30-minute mark with cathode at the right. Scale bar, 200 µm. (C) Hypothesized tug-of-war in cell groups in the presence of EF. The magenta arrow indicates the migration direction for large and small groups and individual cells. The polarization vectors are shown for edge (black) and inner (orange) cells in large and small groups. Note that the edge polarization signals are stronger in the single or edge cell compared to the inner cell.

In the SI, we describe the quantitative details of the model and the search for its mechanical parameters informed by the experimental measurements. Numerical simulations of the model with the constrained parameters reproduced the experimentally observed shapes and movements in four cases of small and large, unperturbed and PI3K-inhibited (Fig. 3B, Movie 6): small PI3K-inhibited groups moved to the anode, in the three other cases, the groups moved to the cathode. The small groups moved faster than the large groups, because the outer cells generate greater net propulsive forces than the inner cells, and in small groups the minority of the inner cells effectively slows the groups down less than the majority of the inner cells in large groups. In addition, because the small groups move faster, lamellipodia at their rears destabilize, further decreasing the resistance to the mechanical majority of the cells and accelerating the groups. This lamellipodia destabilization effect explains the more significant decrease of the group area and increase of the group aspect ratio for the small groups: the outer cells at the rear are not pulled outward against the group direction, allowing the effectively elastic mesh of the inner cells to contract inward, shrinking the rear-to-front distance in the group and decreasing its area. We calculated the displacements, areas and aspect ratios in these four cases, and the results (Table S2) display the same qualitative trends as the experimental measurements (Fig. 3I).

### Bidirectional polarization of complex-shaped group confirms a model prediction

The 2D model also makes a simple prediction about a bilobed-shaped group of PI3K-inhibited cells, with two lobes connected by a narrow cell corridor (Fig. 4A, Movie 7): if the smaller lobe’s size is less than the critical size at which the group directionality switches from the anode to cathode, while the greater lobe’s size is larger than critical, then the lobes would move oppositely to each other in EF, deforming the group shape. We simulated a group of such shape and indeed found that this intuition is correct (Fig. 4A). After such initial bidirectional deformation, the greater lobe prevailed in the tug-of-war and started to drag the whole group to the cathode (Fig. 4A). The cell groups in the experiment tend to stay roundish in the absence of EF, likely because an effective boundary tension generated by a collective myosin contraction of the rears of the outer cells, but, among tens of such groups, we found one large group of PI3K-inhibited cells of the sufficiently elongated shape (Fig. 4B) that allowed us to test this prediction. We observed that in the first 15 min after EF was switched on, all parts of the group moved to the cathode, but the ‘upper’ part of the group less directionally so, than the rest of the group, which pre-deformed the group into the bilobed shape (Movie 8). In the next 15 min, the smaller lobe moved toward the anode, oppositely to the greater lobe that continued to the cathode, as predicted. After that, the cell corridor between the lobes that was narrowing during the bidirectional deformation of the group broke and the two resulting groups went in their opposite respective directions. This last break-up part of the process is not predicted by the model because of the linear elastic character of the cell-cell links in the model, but it is easy to imagine that introducing more complex non-linear dynamic links will lead to accounting for this breakage.

## DISCUSSION

We found that while single unperturbed cells and groups of all sizes move to the cathode, single and small (tens of cells) groups of PI3K-inhibited cells move to the anode, but larger (hundreds of cells) groups still move to the cathode. Groups of the unperturbed cells shift along the EF vector more than PI3K-inhibited cell groups of similar sizes. Smaller groups of the unperturbed / PI3K-inhibited cells are slightly faster than larger groups of the unperturbed / PI3K-inhibited cells, respectively. Large groups of both unperturbed and PI3K-inhibited cells undergo little change in shape and area after initiating the EF-induced motility. When motile, outer cells of such groups had lamellipodia protruding outward all around the group’s periphery. In contrast, small groups of both unperturbed and PI3K-inhibited cells underwent a more significant decrease in area and shape after initiating the EF-induced motility. Such groups also polarized, so that lamellipodia of the outer cells at the front and sides of the small groups were prominent, while those of the outer cells at the rear of the small groups were largely lost, so the small groups’ shapes looked like giant single cells.

We propose the following mechanistic explanation of the data (Fig. 4C): inner cells always, independently of the cells’ chemical state, tend to go to the cathode, while the outer cells behave directionally like single cells. According to this model, unperturbed cells groups of all sizes are driven to the cathode, as are all cells in the group. The directionality of PI3K-inhibited cell groups is determined by a mechanical tug-of-war between the cathode-driven inner and anode-biased outer cells. In larger groups, the former are the majority; they win, and the group migrates to cathode. In smaller groups, the latter are the majority; they win, and the group migrates to the anode. Furthermore, if one additional mechanical assumption is made in the model – lamellipodia of outer cells at the rear of the group, dragged by the group’s majority against their local protrusion directions, are destabilized if the group speed is faster than a threshold – then the model also able to explain semi-quantitatively not only groups’ speeds, but also their shapes and polarization states. In agreement with this model, in rare large PI3K-inhibited cell groups with complex shapes, small and relatively isolated from the main part of the group cell clusters at the groups’ periphery showed tendency to go to the anode, oppositely to the cathodal direction of the main part of the group.

The main premise of our model – that directionalities of the inner and outer cells are different – is supported by several observations of cohesive groups of other cell types [8, 11, 50]. What mechanisms could be behind this difference? It could be based on significant distinctions in cytoskeletal organizations in inner and outer cells [31], the most prominent of which is large lamellipodia of the outer cells (Fig. 1A, 2B,D). Inner cells have only cryptic lamellipodia [49, 51]; active lamellipodia of these cells are suppressed by physical coupling to neighboring cells on all sides [26]. Another distinction is that the outer cells are strongly polarized, while it was shown that myosin distribution in keratocytes inside the cell sheet is unpolarized [31]. (Inhibition of myosin was very informative for deciphering rules of galvanotaxis for single keratocyte cells [30], but not for the cell groups, because upon inhibition of myosin, cohesion between the individual cells in the groups is lost [31,33], therefore, we did not use this perturbation in this study.) Different cytoskeletal structures could interpret extracellular signals in divergent ways [52].

Another possibility stems from different sizes of large lamellipodia of the outer cells and small cryptic lamellipodia of the inner cells [49] and potential signal strength dependence on cell size [29, 42]. Specifically, considering that EF induces a gradient of charged motile proteins across the cell surface [29], the ratio of the charged motile protein concentrations at the cathodal and anodal sides of the lamellipodia is on the order of *exp*(*qEL*/*k*_*BT*_), where *q* is an effective protein net charge, *E* is EF amplitude, *L* is the lamellipodial width, and *k*_*BT*_ is molecular thermal energy. This exponential factor could translate into great size-dependent signal differences, so that larger outer cell lamellipodia perceive a much stronger EF signal that could trigger some intracellular signaling pathways which are not activated by weaker sub-threshold EF signal in smaller inner cells. Notably, small lamellipodial fragments have opposite directionality in EF compared to large whole keratocytes [30]. Lastly, considering the likelihood of several pathways polarizing cells in EF [30], one could hypothesize that the EF signal is transduced by three independent additive pathways. For example, the first pathway weakly polarizes the cell body to the cathode, the second one strongly polarizes the lamellipodium to the cathode in a PI3K-dependent way, and the third one strongly polarizes the lamellipodium to the anode in a PI3K-independent way. (Of note, active PI3K was observed to localize to the leading edge of leader cells, but not in follower cells in the group [50].) Then, in the inner cell, the first pathway becomes dominant orienting the cell to the cathode independent of its chemical state. In the unperturbed outer cell, the effects of the second and third pathways cancel each other, and the first pathway, again, orients the cell to the cathode. Finally, in the PI3K-inhibited outer cell, the second pathway does not work, and the strong third pathway wins against the weak first pathway orienting the cell to the anode.

An interesting open question is how general the size-dependent directionality phenomenon is. One study [53] reported that lymphocyte clusters always followed chemical gradient, independent of its steepness, while single cells reversed directionality for very steep gradients. It is well known in chemotaxis that individual cells can be turned around by some chemical perturbations [54], but how cell groups will behave under the same perturbations is unclear. Another important question is whether there are physiological implications of the reported galvanotaxis phenomenon. When fish skin is injured, keratocyte cell groups migrate to repair the wound and re-epithelialization of these wounds is extremely rapid [55]. Considering that these cell groups follow local EF [18], and that PI3K activity levels can vary in cells engaged in collective locomotion [56], it is to be expected that complex, size-dependent directionality affects wound healing.

There is much complexity of the collective galvanotactic response that our study did not address. There are relatively slow (hours-scale) processes of adaptation of the group migration to EF manifested in gradual slowing down of the migration [31, 33, 57], and in gradual redistribution of global stresses in the groups [26]. Effects of EF on cell motility are more nuanced than orienting and calibrating cell propulsion forces [41]. Other sources of complexity are natural cell-to-cell variabilities in directional responses, nontrivial rheology of the cell clusters and stochastic effects in collective directional sensing [58]. Another complex possibility is that the cells inside the group are integrated into supracellular sheets irreducible to autonomous mechanochemical units [13]. Indeed, integrated cytoskeletal networks [31], intercellular stresses [26] and global bioelectric gradients [59] in migrating cell groups were reported. Last, but not least, there is a subtle interplay between polarization vector and propulsion force in cells, which our model does not take into account; for example, it is possible that main driving force arises from cells inside the group, while outer cells are mainly giving direction [60]. Future research will shed light on the impact of these complexities on collective cell galvanotaxis.

## CONCLUSION

Individual fish keratocyte cells migrate to the cathode, while inhibition of PI3K reverses single cells to the anode. Chemically unperturbed cell groups of any size move to the cathode. Large groups of PI3K-inhibited cells move to the cathode, while small groups of PI3K-inhibited cells move to the anode. These results are most consistent with the model according to which inner cells, whether unperturbed or PI3K-inhibited, always have tendency to move to the cathode, while peripheral cells in the group behave as single cells: they are directed to the cathode if uninhibited, but biased to the anode, if PI3K-inhibited. A mechanical tug-of-war between the inner and edge cells directs larger groups with majority of the inner cells to the cathode, and smaller groups with majority of the outer cells to the anode.

## AUTHOR CONTRIBUTIONS

C.C., Y-H.S., M.Z. and A.M. conceived and directed the study; C.C., Y-H.S. and A.M. wrote the manuscript with help from Y.B.; Y-H.S., C.C., M.Z. and A.M. analyzed the data with help from Y.B.; Y-H.S. did the experiments with help from K.Z., Y.Z. and B.R.; C.C. did the modeling with help from H.Y. and F.L. B.D. provided the cell culture.

## ACKNOWLEDGMENTS

This work was supported by NSF grant DMS2209494 to C.C., US Army Research Office grant W911NF-17-1-0417 to A.M., by Air Force Office of Scientific Research grant FA9550-16-1-0052 to M.Z. (Program PI: Wolfgang Losert), and by National Institutes of Health grant EY 019101 to M.Z.

## FIGURE LEGENDS

**SUPPLEMENTAL FIGURE 1: EF-guided collective cell migration of fish keratocytes**.

A. A Schematic view of the electrotaxis chamber and the setup for EF application.
B. Workflow for image processing. Cell group movement was monitored by overlapped multi-field time lapse recording. Multi-field images were stitched together as described in Methods and assembled into temporally ordered stacks for subsequent quantification.
C. Cell numbers in groups was either counted directly or estimated per squared unit area. Scale bar: 50 μm.
D. Groups were outlined and their movements before and after EF application were tracked manually or automatically in ImageJ. Morphological changes were characterized by shape analysis.

**SUPPLEMENTAL FIGURE 2: Results of the 1D model**.

For the given parameters, group velocity *v* is shown as a function of the number of inner cells *n* along a one-dimensional configuration in the presence of an EF. *v*_*1*_ denotes the uninhibited group velocity when both edge and inner cells exert propulsive forces, while *v*_*3*_ is the same condition but for PI3K-inhibited groups. *v*_*2*_ is the velocity when only the right (cathode-facing) edge cell and inner cells produce propulsive forces, and *v*_*4*_ is the asymmetric velocity for PI3K-inhibited groups. The rupture force of edge cells in uninhibited groups is *v*_*EF*_ (green dashed line) and it is *v*_*LY*_ (gray dashed line) for PI3K-inhibited groups.

**SUPPLEMENTAL FIGURE 3: Lamellipodia at the front and rear of large cell groups**.

A) Phase contrast images of a large keratocyte group at the onset (0 min) and after 15 and 30 minutes of EF application in the indicated orientation (Ctrl). Insets show magnifications of the highlighted cathodal/anodal edge areas, showing intact lamellipodia (dashed enclosures). This group contains approximately 54 cells. Scale bar, 100 μm. Time stamps in mm:ss.

B) Phase contrast images of a large keratocyte group at the onset (0 min) and after 15 and 30 minutes of EF application in the indicated orientation, in the presence of a PI3K inhibitor (LY). Insets show magnifications of the highlighted cathodal/anodal edge areas, showing intact lamellipodia after EF application (dashed enclosures). This group contains approximately 52 cells. Scale bar, 100 μm. Time stamps in mm:ss.

## METHODS

### Primary culture of keratocyte groups and single cells

Adult zebrafish (strain AB), were obtained from the UC Davis Zebrafish facility. All experiments were conducted in accordance with the UC Davis Institutional Animal Use and Care Committee protocol 16478. Scales were removed from zebrafish flanks and allowed to adhere to the bottom of EF chambers [31, 61]. The scales were covered by a glass 22-mm coverslip with a stainless-steel nut on the top and cultured at room temperature in Leibovitz’s L-15 media (Gibco BRL), supplemented with 14.2 mM HEPES pH 7.4, 10% Fetal Bovine Serum (Invitrogen), and 1% antibiotic-antimycotic (Gibco BRL). Scales were removed gently once an epithelial sheet forms, which usually takes 24-48 hours. Cell groups of various sizes and shapes separated from the epithelial sheet spontaneously and migrated away; these groups were used in the experiments with EF. To isolate single cells, groups of keratocytes that migrated off the scale were digested by a brief treatment with 0.25% Trypsin/0.02 EDTA solution (Invitrogen) in phosphate buffered saline (PBS), and then kept on ice until use [61, 62].

### Pharmacological perturbation experiments

Drugs were added in the culture medium in the following concentration: 0.1% DMSO (Sigma, Cat# 900185) as drug control, 50 or 100 μM LY294002 (Sigma, Cat# 440202). Subsequent experiments were implemented in the presence of each of these drugs within 15 minutes of incubation.

### EF application and time-lapse recording

The electric fields were applied as previously described [63, 64] in custom-made electrotaxis chambers to minimize heating during experiment. To eliminate toxic products from the electrodes that might be harmful to cells, agar salt bridges made with 1% agar gel in Steinberg’s salt solution were used to connect silver/silver chloride electrodes in beakers of Steinberg’s salt solution to pools of excess medium at either side of the chamber (Fig. S1A). EF strength was chosen based on physiological range and previous studies [31]. In most experiments, an EF of 1 V/cm was used unless otherwise indicated. The actual voltage is measured by a voltmeter before and after each experiment. Phase contrast images were captured by a Zeiss Observer Z1 (Carl Zeiss) equipped with a QuantEM:512SC EMCCD camera (Photometrics). Time-lapse experiments were performed using MetaMorph NX software controlling a motorized scanning stage (Carl Zeiss). Typically, in each experiment, overlapped fields covering a whole cell group were captured sequentially. Images were taken at 30 or 60 second interval at room temperature for up to 60 minutes unless stated otherwise.

### Image processing and data analysis/presentation

Time-lapse images were imported into ImageJ and stitched by using Grid/Collection Stitching plugin (Fig. S1B). To quantify single-cell and collective group motility, we extracted the trajectory of each object using an automatic/manual tracking tool [31, 66, 67]. Directedness was defined as cos(*θ*), where *θ* is the angle between the EF vector and the vector connecting the centroids of the cell/group in the initial and final positions [38, 65]. To quantify cell/group migration, x and y coordinates of single cells and of the groups’ centroid were measured in each image (the origin (0,0) was the initial coordinate) in the image stack, with the x-axis parallel to the EF, as described previously [31]. The (x, y) trajectory data were imported into ImageJ chemotaxis tool plugin and rescaled to physical dimensions based on pixel sizes. The trajectory speed was computed by dividing incremental centroid displacement over respective time interval and then averaging this momentous speed over the whole trajectory.

### Morphological characterization and quantification

Cell numbers in each group were either manually counted (for the groups with less than 100 cells) using ImageJ particle analysis tool (Fig. S1C, left panels) or calculated based on the area fractions (for the groups with more than 100 cells, Fig. S1C, right panel). Groups were then either manually outlined in ImageJ or processed by using a custom written Matlab codes to outline with extracted contours [68, 69]. Briefly, we used Matlab edge detection and a basic morphology function to outline cell groups in the phase contrast image. In most cases, we had to use the Lasso tool in Photoshop (Adobe) to manually draw the group shape. Polygonal outlines extracted from the binary images were plotted in Celltool, an open source software [70]. Geometric features of each cell group including centroid, aspect ratio, and area (Fig. S1D) were measured directly from the polygons by using standard formulas [53].

### Statistics and reproducibility

All experiments were repeated and produced similar results. In most cases a representative experiment is shown, unless stated otherwise. Data are presented as means ± standard error. To compare group differences (EF vs no EF or drug treatment vs no treatment), either chi-squared test, or paired/unpaired, two-tailed Student’s t-test were used. A p value less than 0.05 was considered as significant.

### Modeling

Details of the computational model are described in the Supplemental information.

## MOVIES

**Movie 1:** Pair of PI3K-inhibited cells without and with EF.

**Movie 2:** Three small groups of unperturbed cells polarizing and initiating collective migration in EF.

**Movie 3:** Four small groups of PI3K-inhibited cells polarizing and initiating collective migration in EF.

**Movie 4:** Large group of unperturbed cells polarizing and initiating collective migration in EF.

**Movie 5:** Large group of PI3K-inhibited cells polarizing and initiating collective migration in EF.

**Movie 6:** 2D simulations of small and large group of unperturbed and PI3K-inhibited cells polarizing and initiating collective migration in EF.

**Movie 7:** 2D simulations of the dumbbell-shaped group of PI3K-inhibited cells polarizing and initiating collective migration in EF.

**Movie 8:** Experimentally observed dumbbell-shaped group of PI3K-inhibited cells polarizing and initiating collective migration in EF.

