## Supplementary figures and images for "Galvanotactic directionality of cell groups depends on group size"

### Supplemental Figure 1

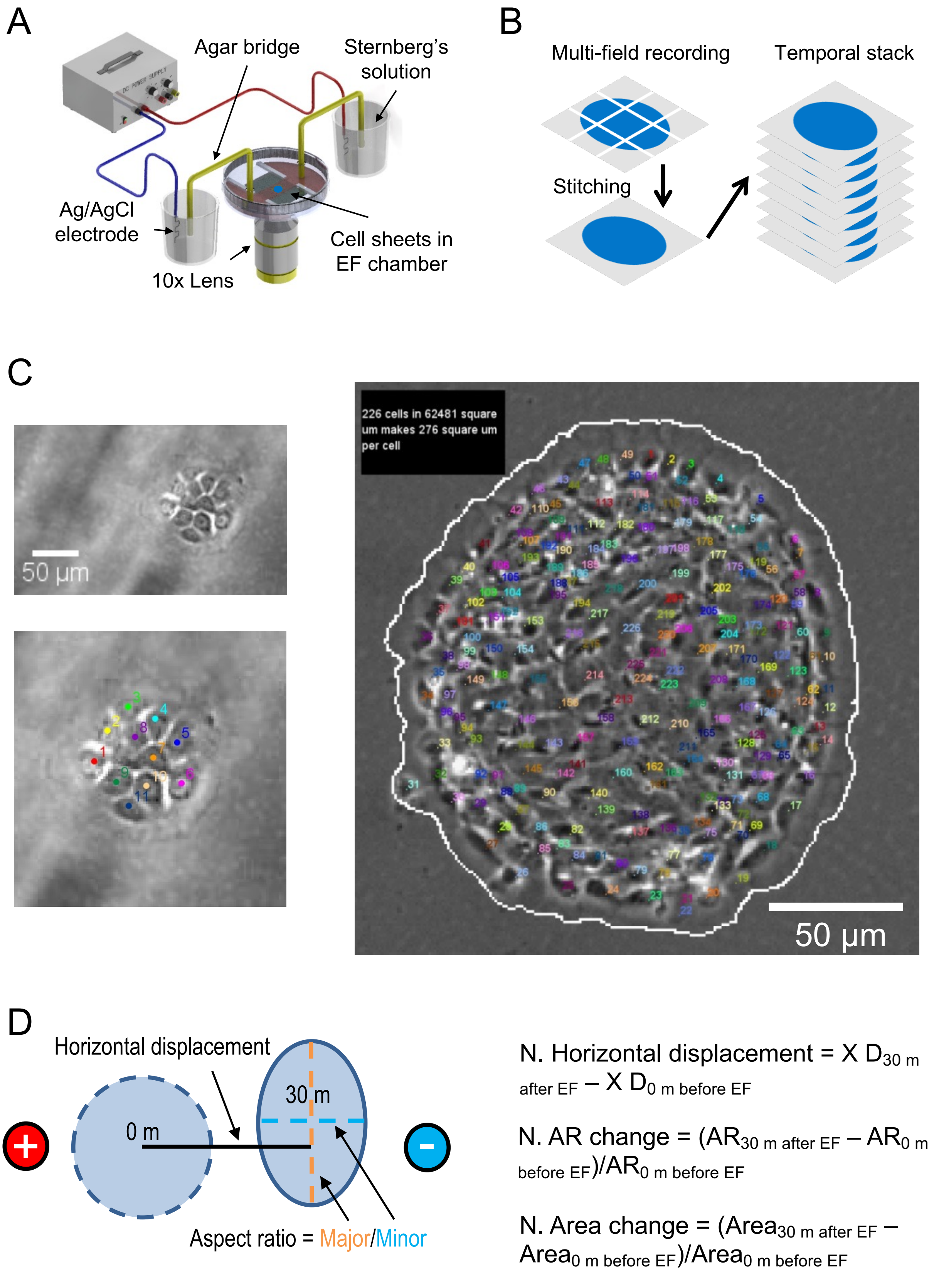

### Supplemental Figure 2

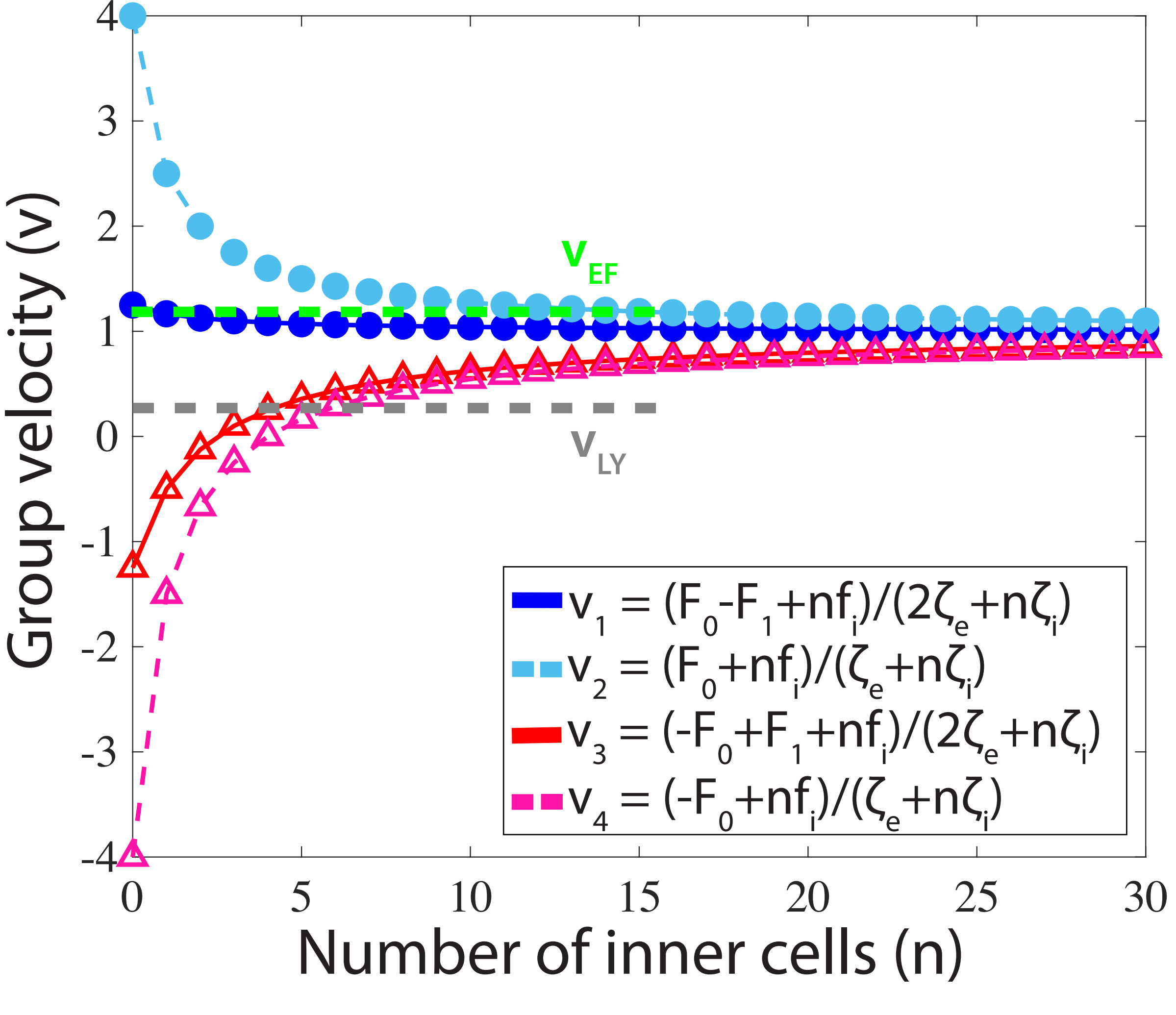

### Supplemental Figure 3

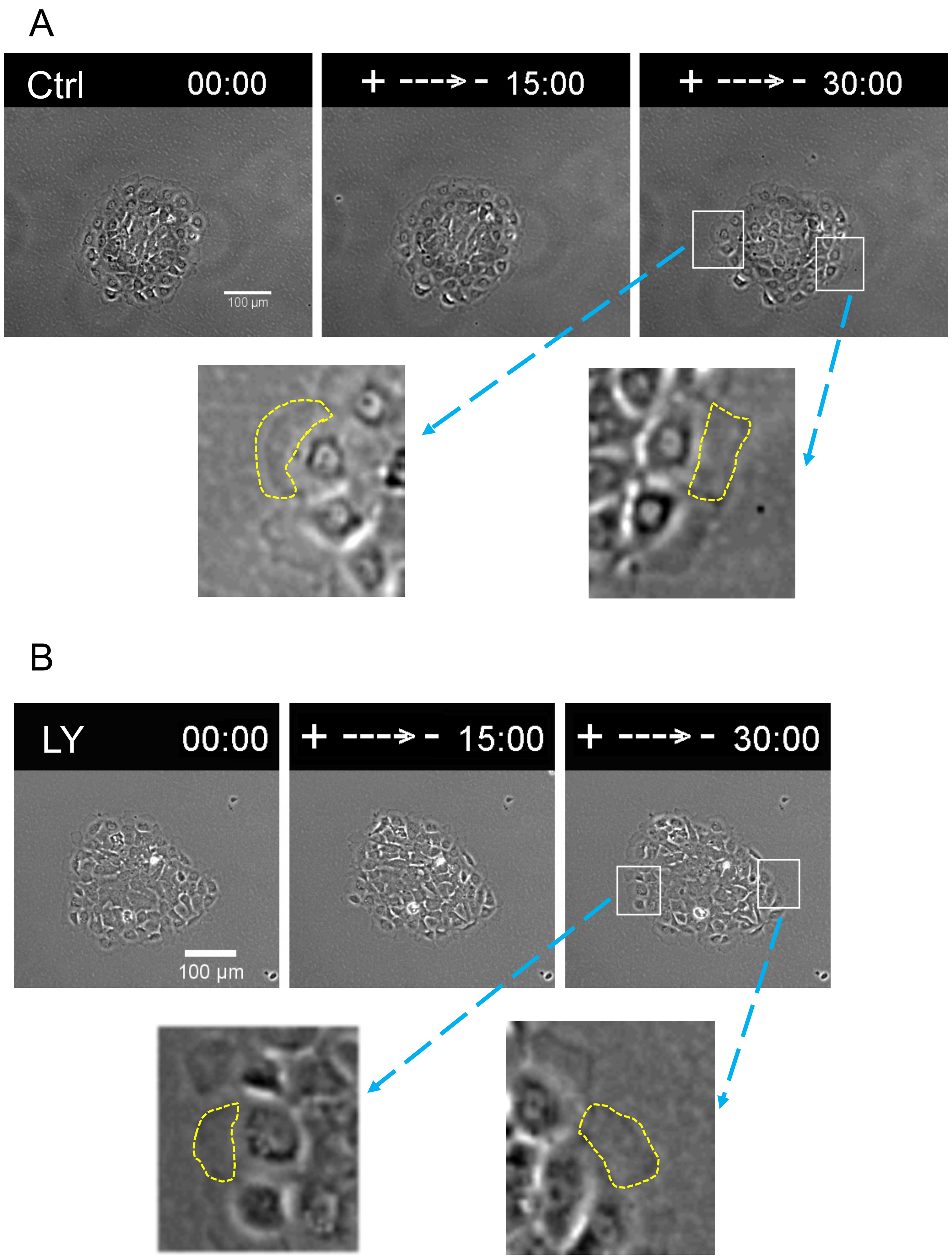


Fig. S3
